# Screening candidate genes related to psoas muscle traits in Debao and Landrace pigs based on transcriptome analysis

**DOI:** 10.1101/2022.04.04.487004

**Authors:** Chang-yi Chen, Su-xian Zeng, Yuan-ding Ma, Jun-wen Zheng, Xin Li, Chen-yong Xiong, Hong-jin Zhou, Chun-tao Wei, Zong-qiang Li

## Abstract

To identify the important genes that affect the phenotypic differences between the Landrace and Debao pigs, especially the differences in metabolism and muscle growth. Differentially expressed genes of psoas major were detected by mRNA transcriptome sequencing in Landrace and Debao pigs. By extracting the total RNA of the psoas major muscle of the Landrace pig and the Debao pig, purifying the mRNA, constructing the cDNA library, conducting transcriptome sequencing, and then through the sequencing quality evaluation, we know that the sequencing quality of this study is relatively high. A total of 17,943 genes were detected in all samples, including 17,870 known genes and 73 new genes. Defined genes with |log2FC| greater than 2 and Q-value less than 0.001, and screened them as significantly differentially expressed genes. A total of 1661 differentially expressed genes were screened from the samples of Landrace pigs and Debao pigs, among which 1255 genes were differentially up-regulated and 406 genes were differentially down-regulated. Through differential gene analysis, it is concluded that these genes are mainly involved in metabolic regulation, muscle and fat development and other processes, especially some important functional genes such as MAPK14, FOS, SIRT1, KRAS, EGR1, CDNNB1, etc. To sum up, this study used transcriptome sequencing method, and then selected differentially expressed genes between Landrace pigs and Debao pigs through data analysis, and finally screened out important genes affecting phenotypic differences, which provided genetic support for breeding better breeds in the future.

## 1. INTRODUCTION

With the development of science and technology, molecular biology has ushered in a new chapter, and a variety of new sequencing technologies emerge in endlessly, such as mRNA transcriptome sequencing, so now part of the research is biased towards the detection of important economic animals [1-3]. With the development of the human genome project, some scientists turn their attention to animals, especially the economically valuable livestock, such as pigs. And because of the maturity of the human genome project, scientists have more experience to study the pig genome. In addition, skeletal muscle is divided into intermediate type, rapid glycolysis type, fast oxidizing type and slow oxidizing type, which have a great relationship with pork meat quality. The psoas major muscle is a very typical slow oxidizing muscle. Transcriptome was first proposed by Velculescu in 1997[4]. Since then, studies on porcine transcriptome have mainly focused on abdominal subcutaneous fat, embryo and longissimus dorsi muscle, etc. Therefore, there are still few studies on psoas major at present, and sequencing data on psoas major transcriptome can provide help for future studies.

In today’s society, people have higher and higher requirements for the quality and taste of pork. More and more people will choose to buy characteristic varieties of pigs at high prices to meet their desire for higher meat taste. The most important factor determining the quality and taste of pork is the fat content of the muscle. Although the feed utilization rate of commercial pigs is high and the growth rate is fast, the fat content in the muscle of these pigs is low, leading to poor taste, such as the Landrace pigs selected in this study. Debao pig, another kind of pig selected in this study, is a unique pig type with high fat content in Debao County, but its growth speed is too slow and feed utilization rate is very low, which can not meet the needs of the society at all. So we chose these two kinds of pigs with obvious phenotypic differences to explore the important genes affecting the differential expression of muscle and fat. But there are many factors that affect the meat quality and taste of pork, not only muscle and fat, so we need to explore a variety of metabolic pathways, such as differential expression of these pathways and phenotypic differences. To sum up, we will provide genetic support for breeding excellent pig breeds in the future by sequencing the whole gene transcriptome of Landrace and Debao pigs and analyzing the differential gene expression.

## 2. Materials and methods

### 2.1 Experimental materials

Three Landrace pigs and three Debao pigs were selected from Debao County, Guangxi Province. The six selected pigs were individually raised in separate stalls from birth to 7 months of age, with free access to food, water and both breeds were fed the same conventional diet.Before every swine was slaughtered, a warm shower to relax the pigs, then they were stunned with the low-voltage electric shock to reduce the pain and were exsanguinated by puncturing carotid artery to death, and 20g sample of the Psoas major was collected.The water stains were sucked dry and cut into small pieces (about 1g, thickness less than 0.5cm), and then put into the marked 2.0ml EP tube. After rapid liquid nitrogen freezing, the EP tube was sealed with sealing film, and then placed in a 50ml centrifuge tube or sealing bag, After being transported back to the laboratory, it was stored in the refrigerator at -80°C, and the total RNA was extracted in the laboratory. All animal procedures were approved by the Committee on the Ethics of Animal Experiments of Guangxi University (Protocol Number: GXU2017-014) and were conducted in accordance with the National Research Council Guide for Care and Use of Laboratory Animals (2017).

### 2.2 Experimental processes

Six samples were pretreated first, and then the total RNA was extracted from the samples. In addition to mRNA, ncRNA also exists in the total RNA of eukaryotes, so the total RNA needs to be enriched. The mRNA fragment was used as the template to synthesize single strand cDNA, and then double strand cDNA was synthesized. After synthesis, the double strand cDNA was purified and recovered, the end was repaired, and the junction was connected. Then the fragment size was selected, and then the cDNA library was constructed by PCR amplification. The fragment size and concentration of the library were detected by biological analyzer. After cyclization of PCR products and elimination of non cyclized DNA, the final library was obtained and sequenced. Before the experiment, sterilize the equipment required by autoclave; after the reagents are prepared, put them into the super clean table or refrigerator for use. Low temperature centrifuge is pre-cooled to 4°C in advance for use. Total RNA was extracted by Trizol method, and the concentration, total amount, RIN value and 28S/18S of total RNA were detected by Fragment Analyzer.

### 2.3 Data analyses

The data obtained by sequencing are the original data, and then the quality control of the original data is needed. Clean reads obtained were compared. After that, the second quality control was carried out. Then, the analysis of gene expression level was done, and differentially expressed genes among samples were selected for enrichment analysis, cluster analysis and protein interaction analysis. And then does the quality control.

### 2.4 Difference analyses

The differences of samples were analyzed by principal component analysis (PCA) and correlation analysis. Used to help understand a particular gene set in a certain metabolic pathways, molecular function or participate in biology are significant in the process of enrichment. The specific gene sets mentioned here can be differentially expressed genes, differentially, expressed miRNA target genes, circRNA-derived genes and other gene sets that need special attention. These gene sets are collectively referred to as candidate genes for convenience of expression. In smallRNA analysis, candidate genes used for enrichment analysis refer to target genes with differentially expressed miRNA, and candidate genes refer to circRNA source genes in circRNA analysis. In other RNA analyses, candidate genes refer to differentially expressed genes.

In order to improve the accuracy of DEGs, we define |log2FC| greater than 2 and Q - value of 0.001 or fewer genes, screening for significantly differentially expressed genes. Then the differentially expressed genes were mapped to each term in the database to obtain the gene number of each term. Then, through the test, GO items that were significantly enriched compared with all gene backgrounds of the species were selected from the differentially expressed genes for GO enrichment analysis. Then do KEGG pathway enrichment analysis and the analysis of protein interaction network

## 3. RESULTS

### 3.1 Total RNA quality tests

From the quality test results, we can know that on the one hand, Although the 28s / 18S values of CB_ PM2、CB_ PM3、DB_ PM1、DB_ PM2 and DB_PM3 samples were all less than the standard value of 1.5, the RIN values of PM3 samples were not less than 8, which can ensure the quality of transcriptome library; on the other hand, although the CB values were lower than the standard value of 1.5, the RIN values of PM3 samples were not less than 8_ The 28S / 18S value of PM1 sample is less than the specified standard value of 1.5, and the RIN value of this sample is less than 8, which belongs to risk sample, but the sample base is flat. Therefore, in conclusion, the total RNA quality of these six samples meets the requirements, and can be used for transcriptome sequencing. The results of total RNA quality test are shown in Table 1.

**Table 1.**
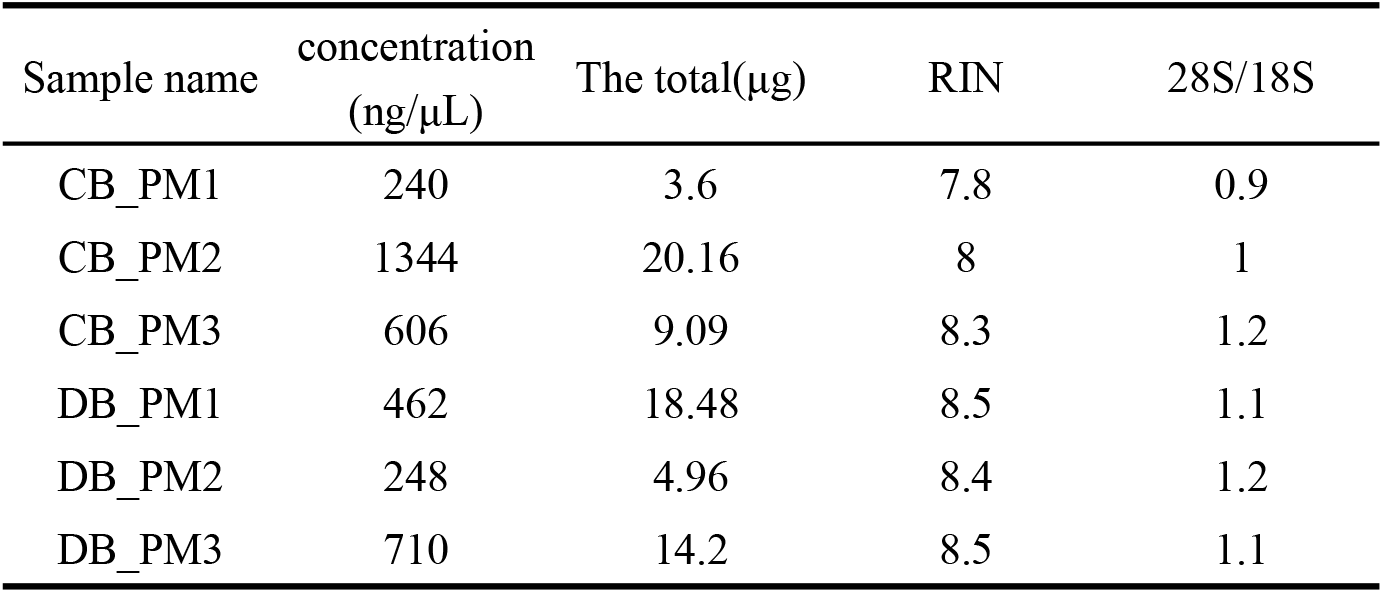
Total RNA quality test results

By analyzing the randomness, saturation and coverage of sequencing, it is known that the sequencing quality is good

### 3.2 Gene statistics

#### Number of sample genes

The average comparison rate of the sample comparison genome was 92.71%, and the average comparison rate of the comparison gene set was 64.52%; a total of 17,943 genes were detected, the number of known genes was 17,870, and the number of new genes was 73.

#### Statistics of various types of RNA

The sample counts the number of various types of RNA through detection. As showed in Figure 1, the number of mRNA measured is 80678, accounting for 78.13% of the total RNA number, and the number of miRNA measured is 457, accounting for 0.44 of the total RNA number. %, the number of lncRNA was 22126, accounting for 21.43% of the total RNA.

**Fig.1.**
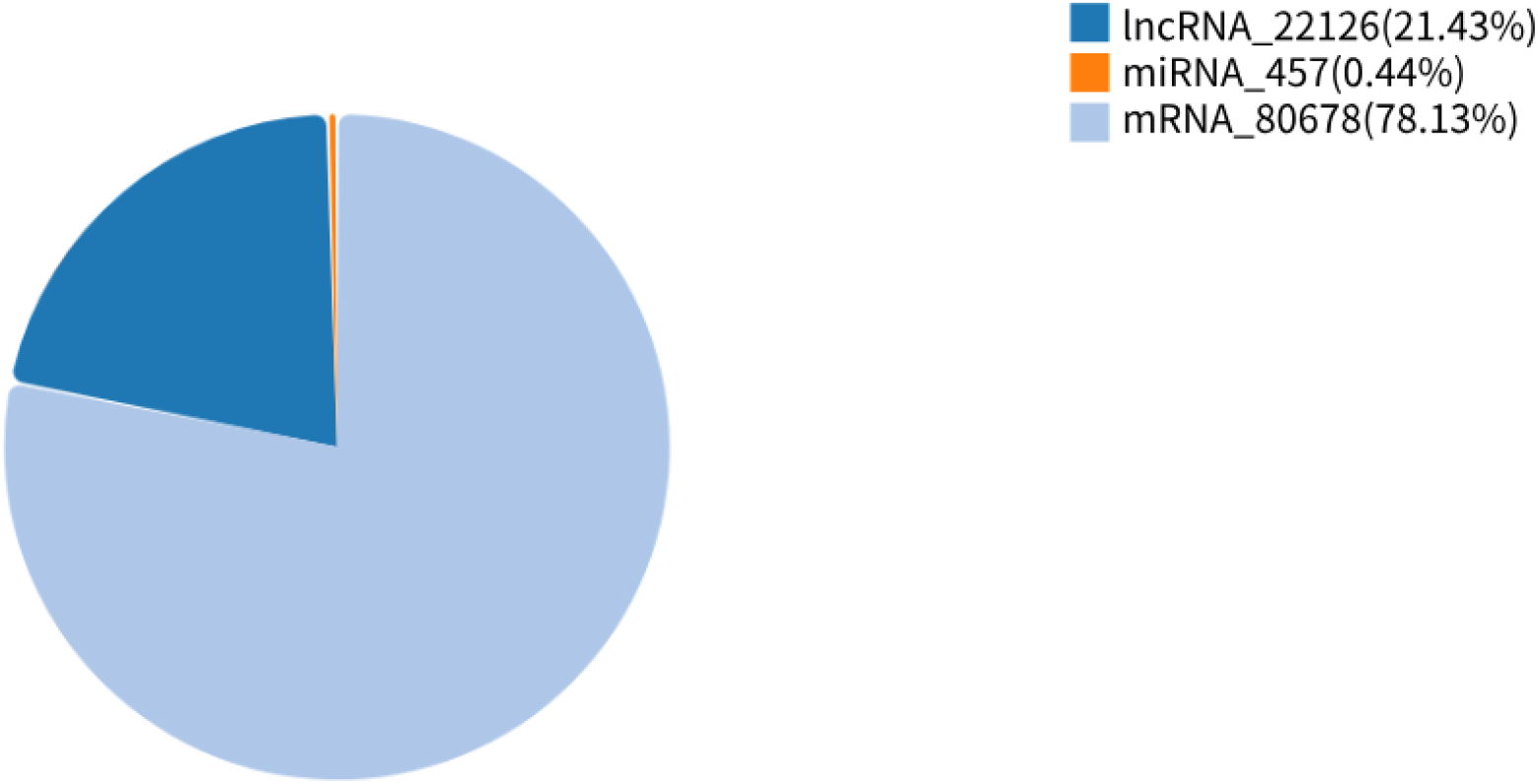
The number of each type of RNA was counted

### 3.3 Difference analyses

#### Principal component analysis

Through principal component analysis of gene expression in 6 samples of Landrace pigs and Debao pigs, the results are shown in Figure 2. We can conclude that the main components of the gene expression levels of samples of the same breed of Landrace and Debao pigs are similar.

**Fig.2.**
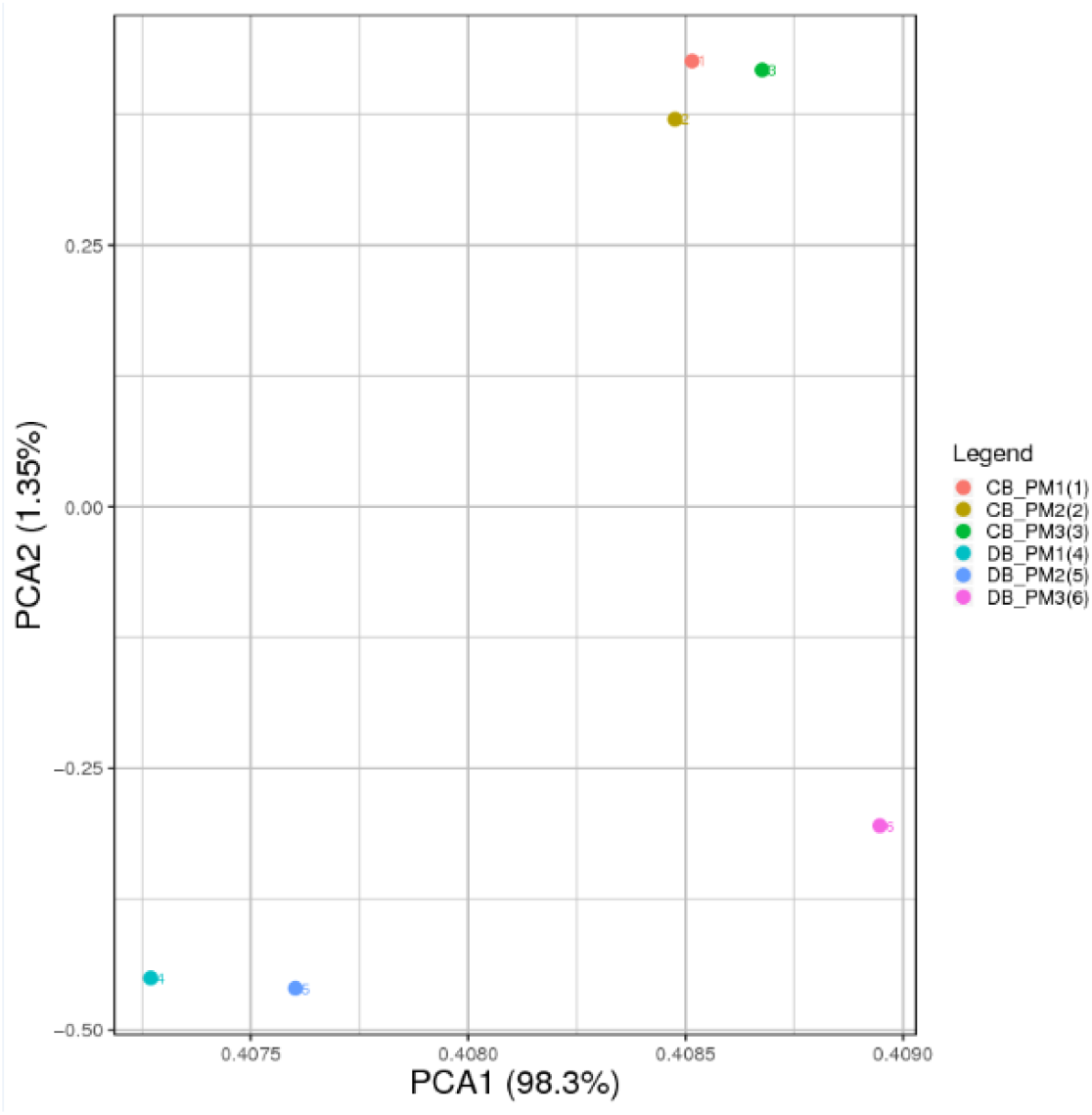
PCA results of samples

#### Sample correlation heat map

Plot the correlation between Landrace pig and Debao pig gene expression, as showed in Figure 3. From the figure, we can know the average Pearson coefficient between Landrace and Debao pigs is 0.9697, which shows that the correlation coefficient between the two samples is large and the sample repeatability is good.

**Fig.3.**
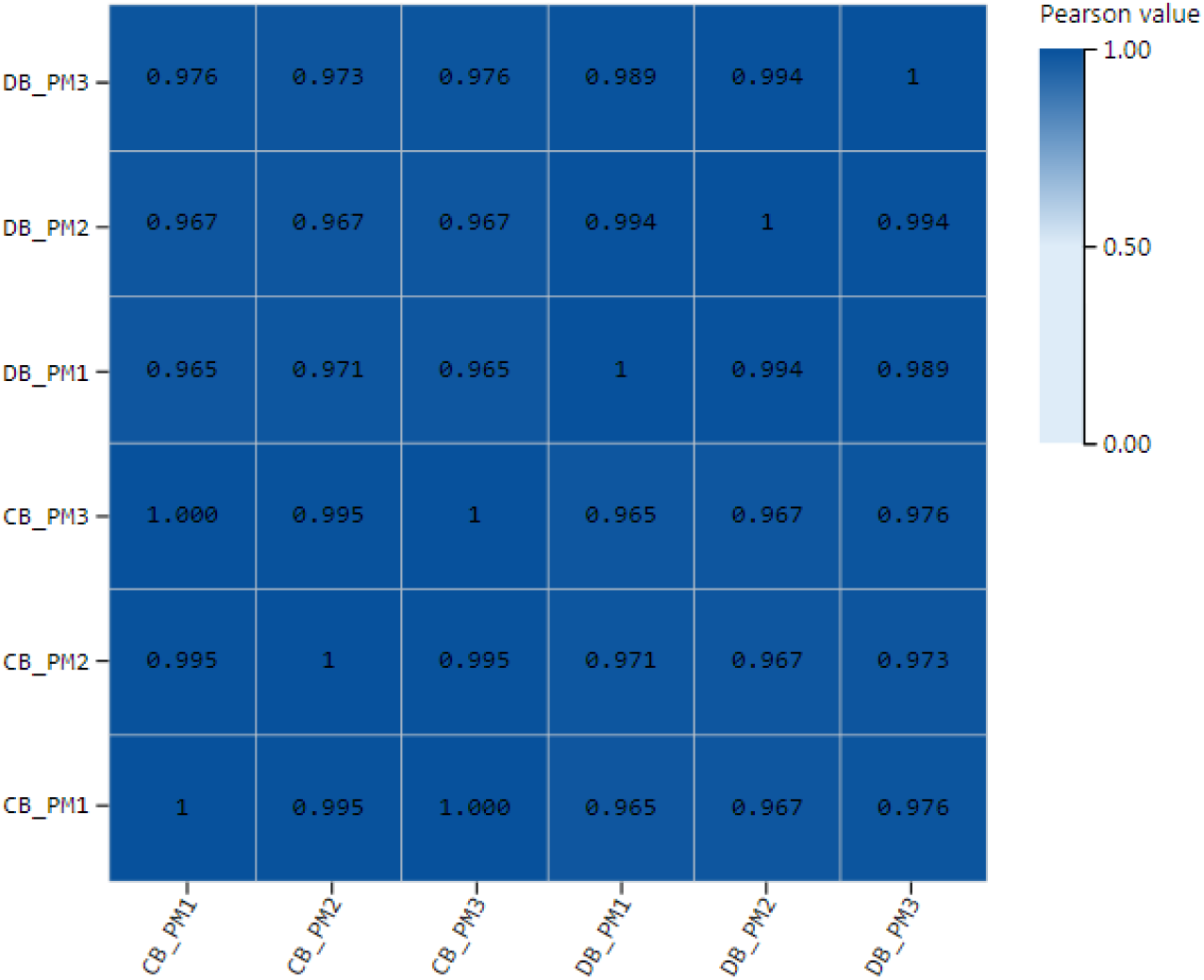
Landrace and Debao pig correlation char

### 3.4 Analysis of differentially expressed genes between samples

#### Statistics of differentially expressed genes between samples

We used statistics of differentially expressed genes between samples of Landrace pig and Debao pig psoas major, and used volcano graphs to show this difference, to show significantly up-regulated, down-regulated, and non-significantly differently expressed genes. As showed in Figure 4.

**Fig.4.**
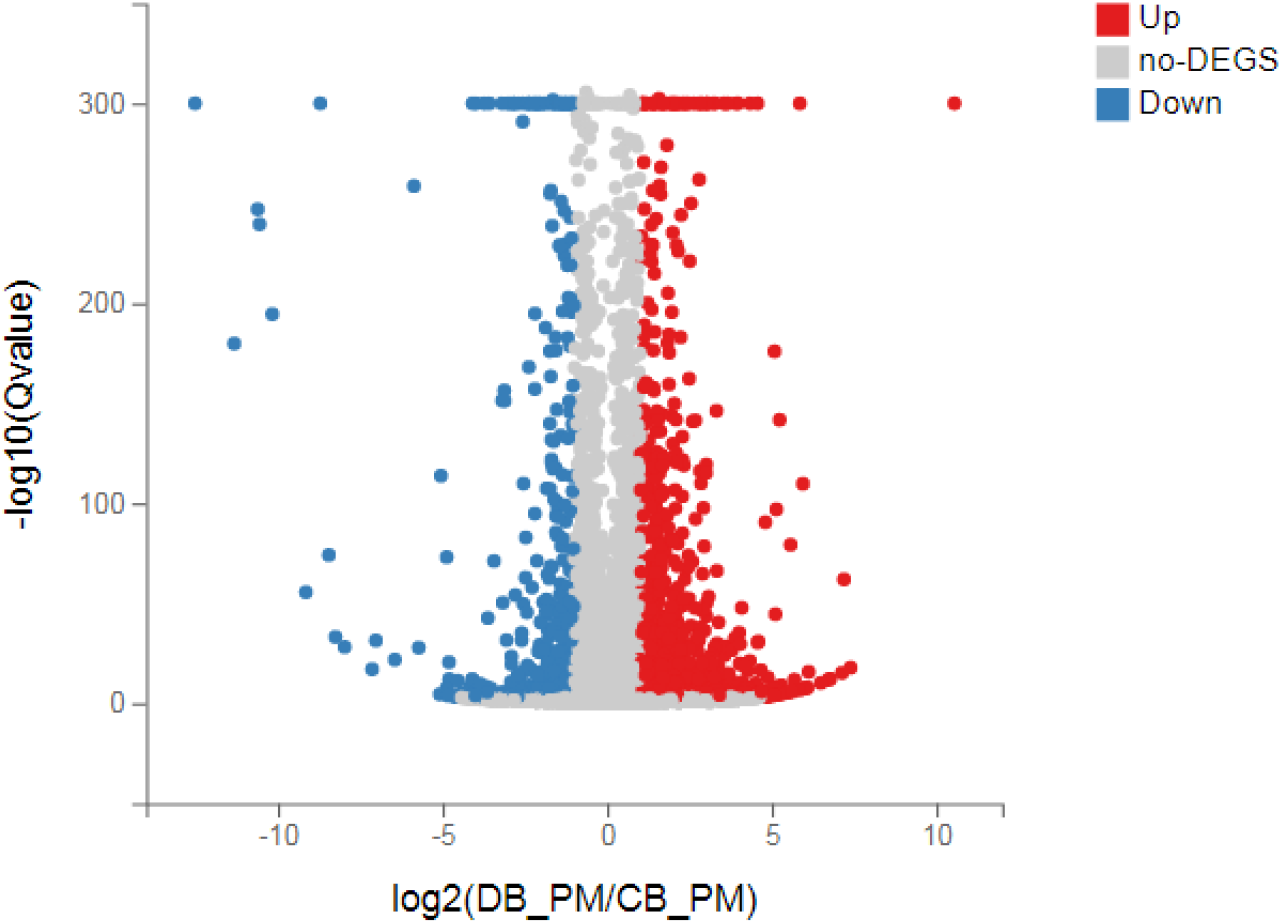
Differential gene volcano plot

Through BGISEQ, we found that the psoas major of Landrace pig and Debao pig had a total of 1661 significantly differentially expressed genes, of which 1255 differential genes were significantly up-regulated and 406 differential genes were significantly down-regulated.

#### Cluster analysis of differentially expressed genes

Through the analysis results of gene expression and gene differential expression, the gene clustering results are intuitively explained through heat maps, and the relationship between differentially expressed genes is clustered. The gene expression of each sample is calculated using logarithm as the base 2, and then the cluster analysis of the detected genes is completed. In the figure, each data corresponds to a row in each figure, and the color is used to determine explain the amount of gene expression. The higher the expression, the red and darker the color. On the contrary, the bluer the color, the lower the expression. The results of cluster analysis of gene expression patterns of all samples are shown in Figure5. We found that the closer sample clustering relationship between the same pig breeds indicates that the sample repeatability is better, and most of the gene expression patterns between the two pig breeds are very different, with opposite trends. Some genes are expressed at a higher level in Debao pigs and at the same time expressed at a lower level in Landrace pigs, and vice versa. Clustering the expression value of the differential gene in each comparison group, the horizontal axis is the log2 (expression value + 1) of the sample, and the vertical axis is the gene. In the default color scheme, the redder the color of the color block, the higher the expression, and the bluer the color, the lower the expression.

**Fig.5.**
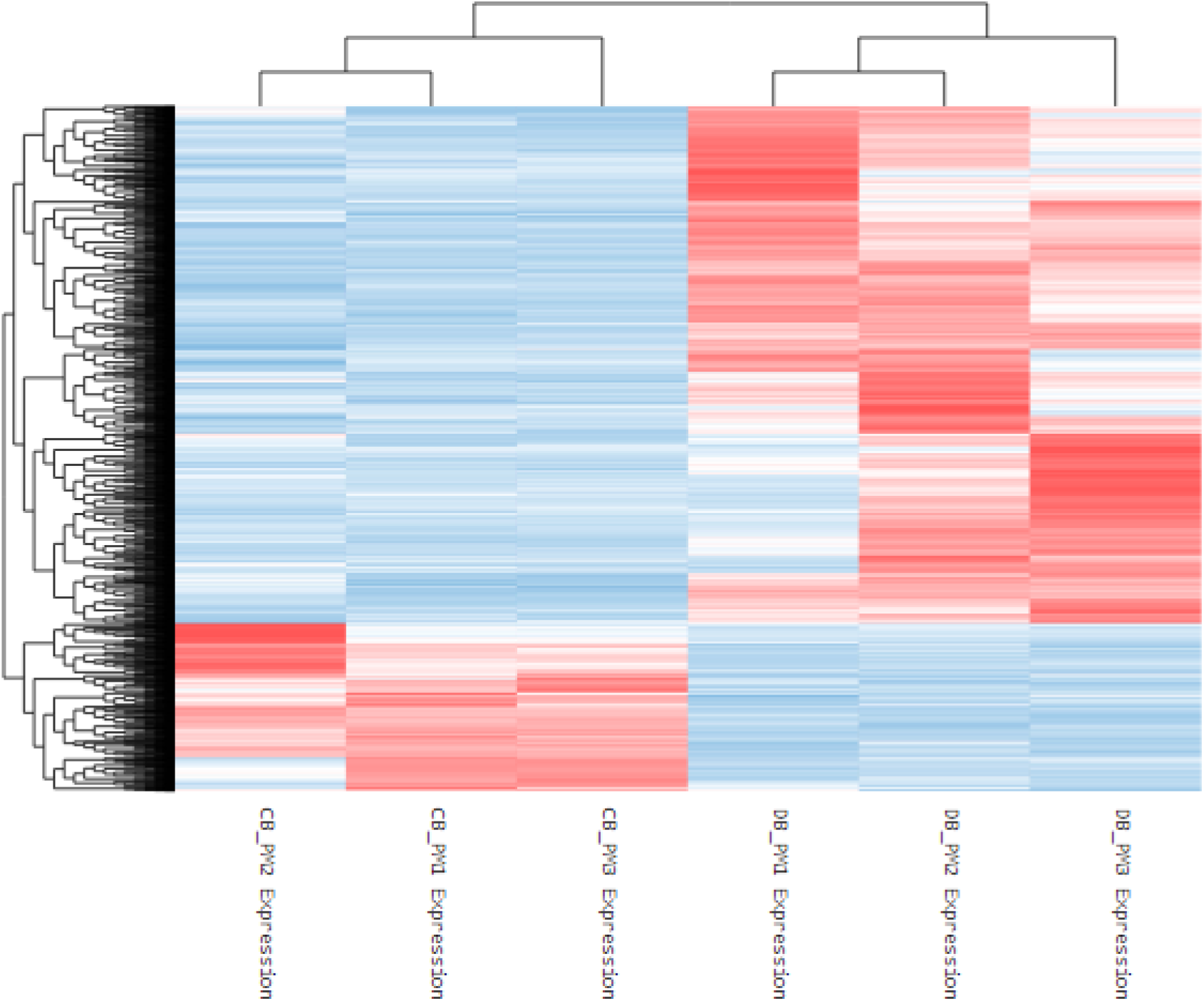
Express quantity is clustering heat map

#### Functional enrichment analysis of differentially expressed genes

In order to better understand the function of the selected differentially expressed genes and the relationship with the phenotype, we carried out a purposeful preliminary screening of the 1661 differentially expressed genes, that is, to screen out the functions and metabolism, tissue and organ development, Related to muscle tissue and structural development, 282 differentially expressed genes were screened, and then functional enrichment analysis was performed, including GO and KEGG signal pathway analysis. We also perform GO term classification statistics for differential genes. Including molecular function, cellular component, and biological process involved, we found that 282 differentially expressed genes can be enriched into 3284, 495, and 335 GO terms in three categories. After screening out Qvalue≤0.05, they are 1608, 181, and 79 respectively. Since a gene often has multiple different functions, the same gene will appear under different classification items. We have shown each of the top 20 items, as shown in Figure 6(a), (b) and (c).

**Fig.6(a).**
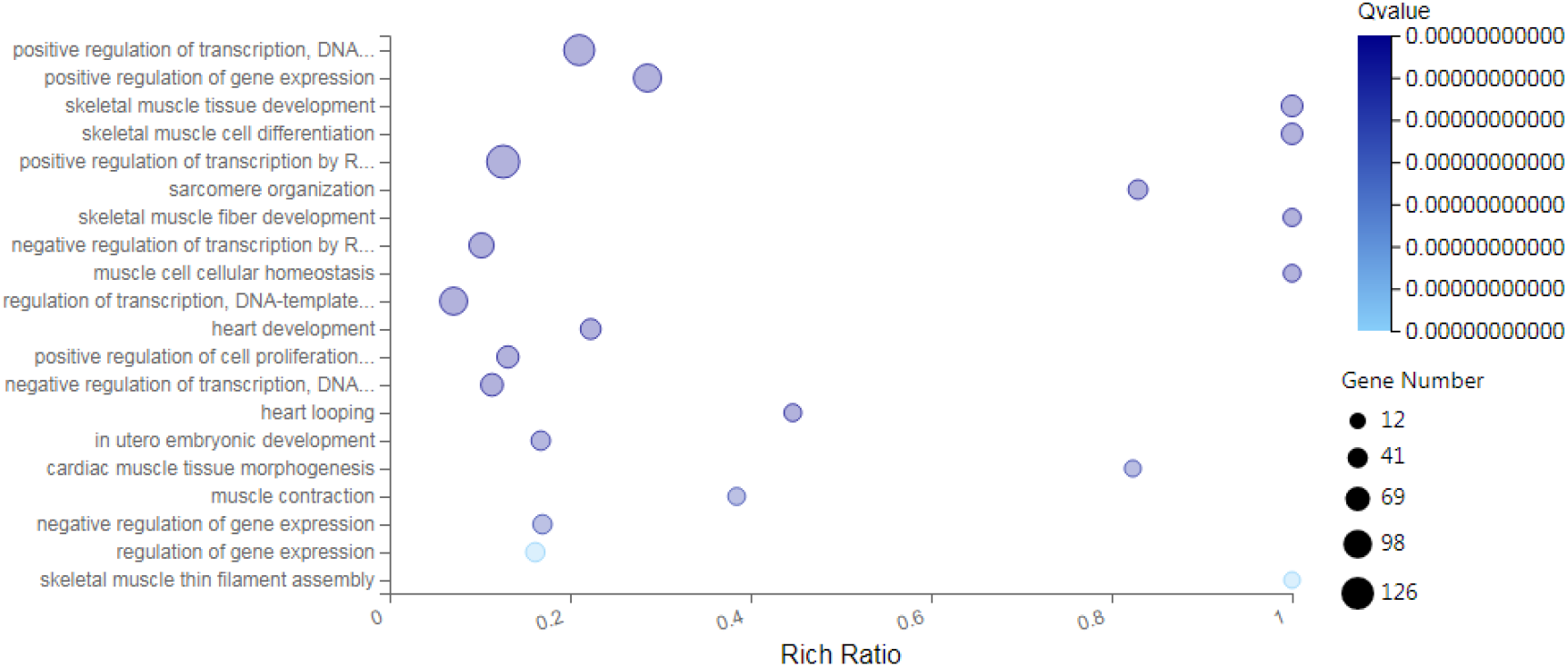
GO analysis of differential genes in biological process

**Fig.6(b).**
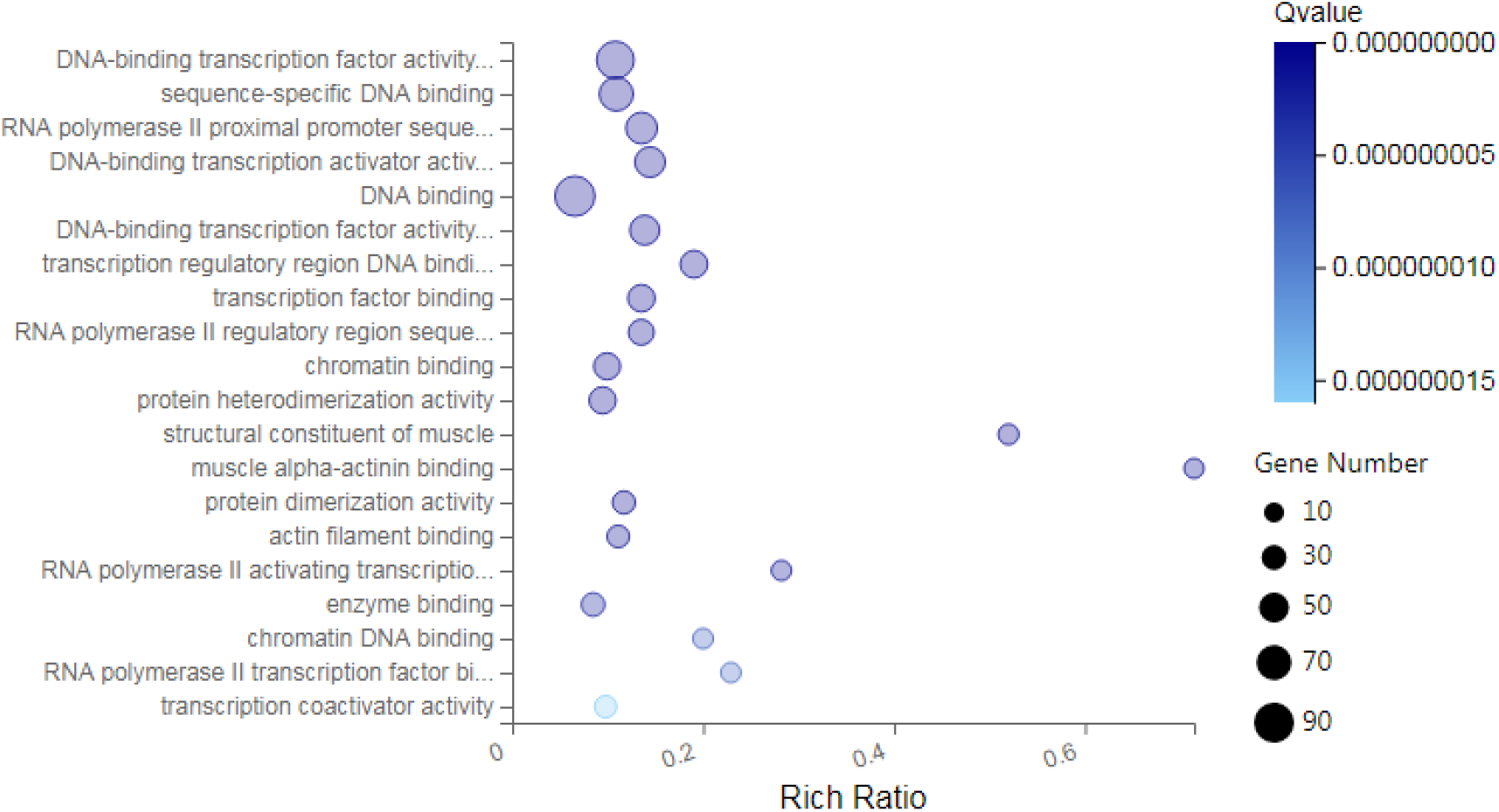
GO analysis of differential genes in molecular function

**Fig.6(c).**
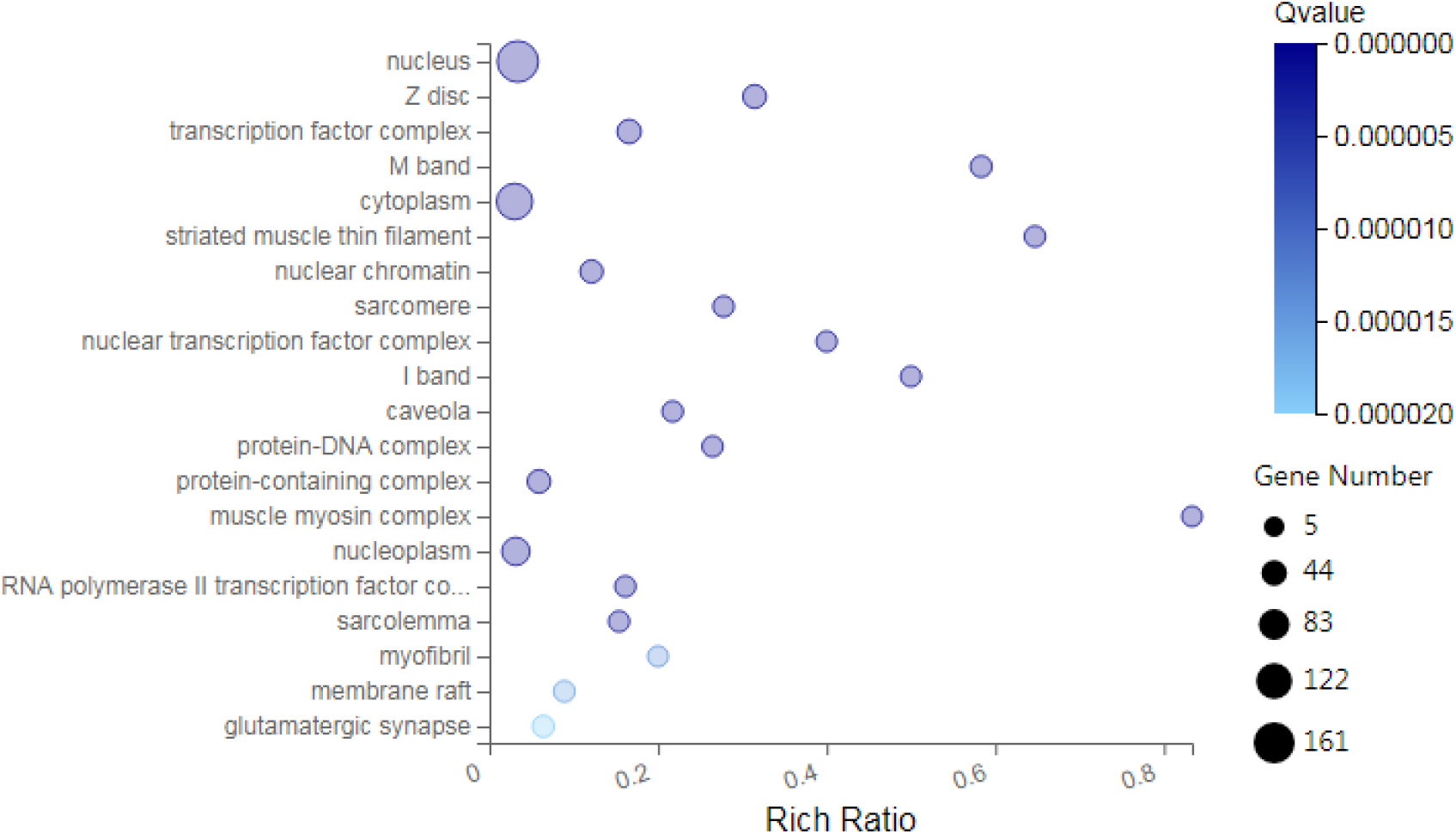
GO analysis of differential genes in cellular component

We found that the function and metabolism of many differentially expressed genes are related to the development of tissues and organs, muscle tissue, and structural development. For example, the genes AKR1C4, MTM1, PTGIS, DNMT1, LOC102167481, GAA, LOC100521659, CBR2, PTGDS, PRXL2B, LARGE1, AKR1C1, ALOX15, LOC100739101, CBR3, LOC102165015MY, ALDH1C2 etc. are involved in the metabolism process, the genes MYL6B, FOXP2, MAPK14, EGR2 and FOS are involved in muscle tissue development.

In order to have a deeper understanding of the functional differential gene expression of the psoas muscle of Landrace and Debao pigs, then we conducted enrichment analysis of the KEGG signaling pathway for these differentially expressed genes. One gene can participate in multiple KEGG signaling pathways. We found that these genes are involved in a total of 166 KEGG signaling pathways. After screening for Qvalue≤0.05, they participated in 70 KEGG signal pathways, we have shown the first 20 signal pathways as showed in Figure 7.

**Figure 7.**
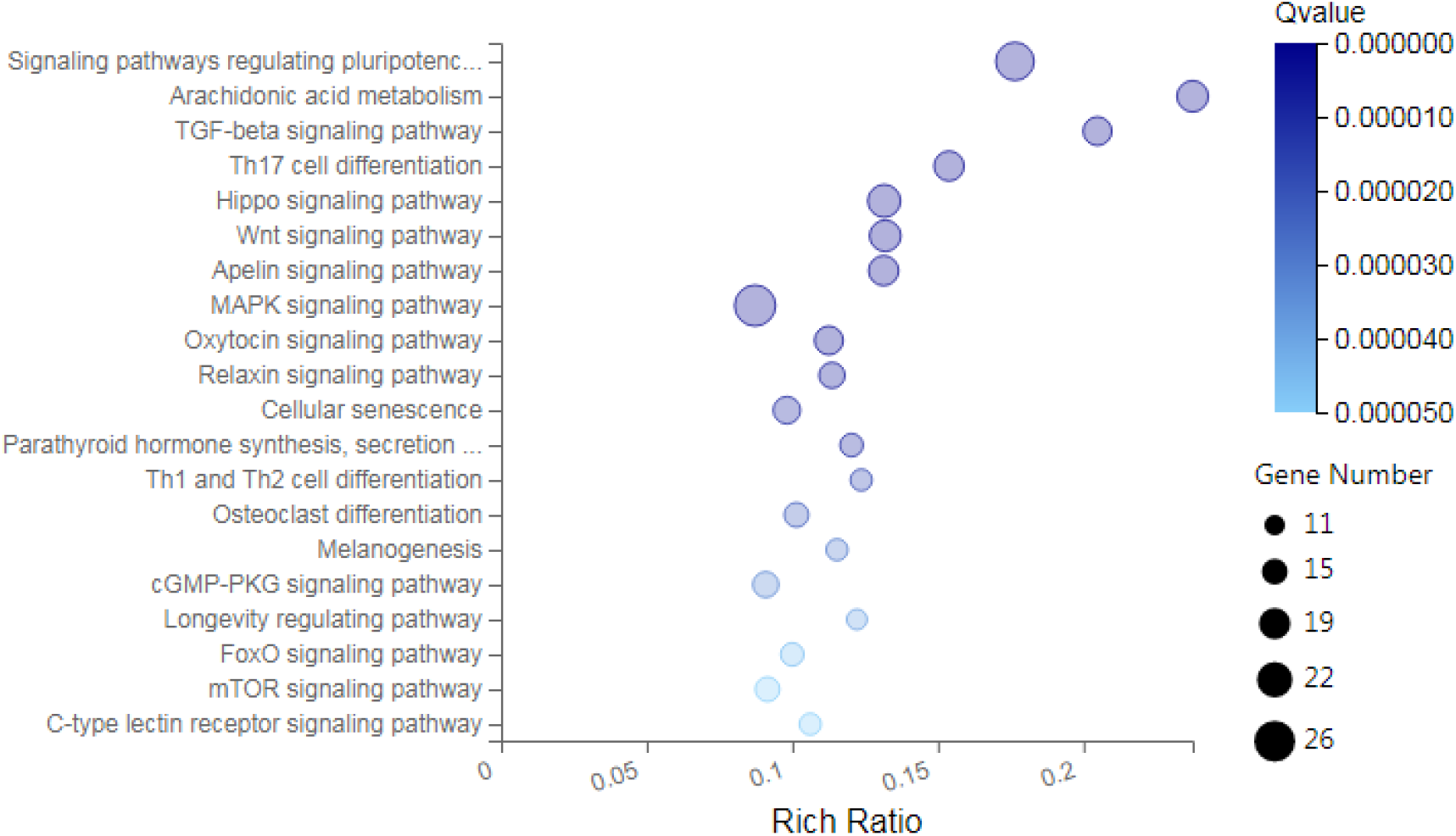
KEGG gene signals pathway enrichment analysis

It is worth noting that there are 18 genes in the number of genes in metabolic pathways, such as: AKR1C4, MTM1, PTGIS, DNMT1, LOC102167481, GAA LOC100521659, CBR2, PTGDS, PRXL2B, LARGE1, AKR1C1, ALOX15, LOC100739101, CBR3, LOC102165015, ALDH1A2, AKR1C2.

### 3.5 Protein network interaction analyses

In order to find the key genes, the online database BGISEQ was used to analyze the protein network interaction of the selected differentially expressed genes. 282 differentially expressed genes were selected through enrichment analysis, and then BGISEQ was used to make a protein network interaction map, as showed in Figure 8.The red-purple dots indicate genes with node connection greater than 10. We obtained 117 nodes in total. Genes with node connection greater than 10 are key genes, such as MAPK14, EP300, SIRT1, KRAS, FOS, and EGR1. These genes are related to metabolic signaling pathways and muscle development.

**Fig.8.**
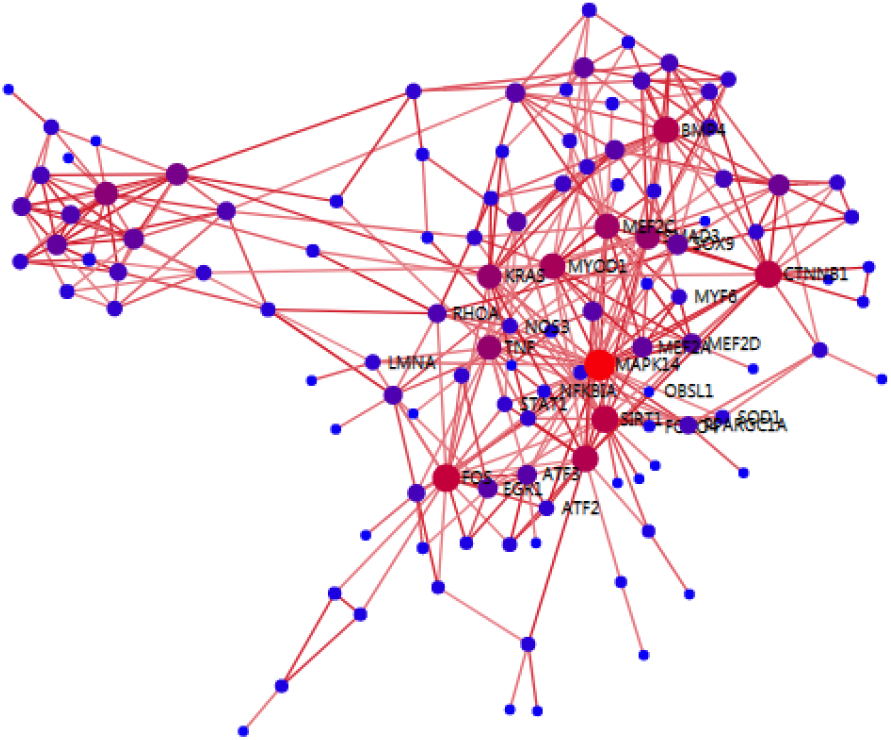
Protein network interaction analysis

## 4. Discuss

At present, many related researches on pigs are aimed at improving the meat quality and taste of pigs. Skugor, A et al. [5] studied the effect of long-term feeding rapeseed powder on pig meat quality and shape, and many studies improved pig breeds to improve pig meat quality [6-9]. In recent years, many studies are based on RNA sequencing technology and differential expression analysis [10-13]. There are many factors that affect the muscle and fat of pigs, such as the influence of gender on the meat quality of pork [14-15]. In this experiment, Landrace pigs which are high feed utilization rate, fast growth rate, but low muscle fat content, poor taste [16] and Debao pigs which are high fat content, but slow growth rate, low fat content, good taste were used to explore the important genes affecting the differential expression of muscle and fat.

In this study, the total RNA of the psoas muscle of Landrace and Debao pigs was extracted, and mRNA was purified to construct a cDNA library and perform transcriptome sequencing[17-19]. The quality of cleaning reads obtained in this study is good. Through gene statistics, we can get that the average comparison rate of the sample comparison genome is 92.71%, and the average comparison rate of the comparison gene set is 64.52%; a total of 17,943 genes have been detected, and the number of known genes is 17,870. The number of new genes is 73.

The distribution of expression levels can help us select genes with high expression levels. Through the statistics of alternative splicing events, we can see statistics of 5 kinds of alternative splicing events, which lay the foundation for gene fusion, which can provide help for cancer research in the future. Through difference analysis, principal component analysis and correlation heat map analysis, we can get that the average correlation coefficient of the two pig genes is 0.9697, indicating that the gene correlation coefficient of Landrace pig and Debao pig psoas major is large and the sample is repeatable it is good.

In recent years, Zhang Xiaoli and others have performed transcriptome sequencing on the longissimus dorsi and psoas major of Landrace pigs, and screened the differentially expressed genes in the longissimus dorsi and psoas major of Landrace pigs through bioinformatics analysis, and obtained 56 up-regulated genes and 137 down-regulated genes. In this study, through analysis and statistics of differentially expressed genes between samples, we can see that there are a total of 1661 differentially expressed genes in Landrace pigs and Debao pigs, of which 1255 are significantly up-regulated and 406 are significantly down-regulated. Finally, through enrichment analysis and protein-protein network interaction analysis, we can select many important genes that affect the phenotypic differences between Landrace pigs and Debao pigs, and the functions of some differentially expressed genes are related to the synthesis of muscle and fat. There are also some differentially expressed genes related to important metabolic signaling pathways. Metabolic regulation and muscle development play an important role in the growth and phenotype of pigs. For example, AKR1C4 MTM1 PTGIS DNMT1 LOC102167481 GAA LOC100521659 CBR2 PTGDS PRXL2B LARGE1 AKR1C1 ALOX15 LOC100739101 CBR3 LOC102165015 ALDH1A2 AKR1C2 and other genes are involved in the development of metabolic signaling pathways, such as MAPK14, FOS, SIRT1, EGR1 and other genes involved in the development of muscle. FOS is an inflammatory and apoptosis-related gene[20-21], and in recent years, studies have reported that FOS may be related to pig muscle growth weight [22]. Studies have also shown that EGR1 is an early growth response protein, which has an important relationship with cell proliferation, differentiation and apoptosis, Egr1 expression is activated by a variety of MAPK-inducing factors[23-25], Multiple studies have shown that the MAPK pathway regulates the expression of adipogenesis transcription factors during adipogenesis [26-27]. EGR1 knockout in mice can reduce leptin expression in basal cells [28]. Studies have also found that EGR1 has an important relationship with the content of muscle fat [29]. The expression of SIRT1 in satellite cells is important for the functional development of skeletal muscle both initially and for full muscle regeneration[30], and recent studies demonstrated that SIRT1 plays a central beneficial role in controlling hepatic lipid metabolism[31]. UCP3 is a negative regulator of energy balance and lipid metabolism. UCP3 gene can affect lipid metabolism and inhibit fat deposition by inhibiting the accumulation of triglycerides in storage cytoplasm [32-34]. This provides genetic support for breeding better varieties in the future.

## 5. Conclusion

This study compared the transcriptome sequencing results of psoas major muscle of Landrace pigs and Debao pigs, and the sequencing quality assessment results showed that the sequencing quality of this study was high and the sequencing data volume met the requirements. A total of 17,943 genes were detected in all samples, including 17,870 known genes and 73 new genes. A total of 1661 differentially expressed genes were screened between Landrace pigs and Debao pigs, among which 1255 differentially up-regulated genes and 406 differentially down-regulated genes were detected. At the same time, the analysis of differential genes showed that these genes were mainly involved in metabolic regulation, muscle and fat development and other processes, especially some important functional genes such as MAPK14, FOS, SIRT1, KRAS, EGR1, CDNNB1, UCP3 etc..

## Acknowledgments

We thank Guangxi Baise Paiqi Co., Ltd., China, for providing the samples during the experiment

## Availability of data and materials section

The datasets generated and analysed during the current study are not publicly available due to the data relate to ongoing post study but are available from the corresponding author on reasonable request.

## Ethics declaration

### Ethics approval and consent to participate

All animal procedures were approved by the Committee on the Ethics of Animal Experiments of Guangxi University (Protocol Number: GXU2017-014) and were conducted in accordance with the National Research Council Guide for Care and Use of Laboratory Animals (2017). All methods were carried out in accordance with relevant guidelines and regulations. This study was carried out in compliance with the ARRIVE guidelines. Before every swine was slaughtered, a warm shower to relax the pigs, then they were stunned with the low-voltage electric shock to reduce the pain and were exsanguinated by puncturing carotid artery to death

### Author Contribution

C.-Y.C., Y.-D.M. and Z.-Q.L. drafted the manuscript. C.-Y.C. and X.L. developed and framed the research questions. C.-Y.C., S.-X.Z. and Y.-D.M analyzed transcriptomics data. J.-W.Z., H.-J.Z., C.-T.W. and C.-Y.X. contributed to serum biochemical analysis and subsequent data analysis. Y.-D.M. and X.L. contributed to revision of the entire manuscript. All authors have read and approved the manuscript.

## References

[1] Drag, M., Hansen, M. B., & Kadarmideen, H. N. (2018). Systems genomics study reveals expression quantitative trait loci, regulator genes and pathways associated with boar taint in pigs. PloS one, 13(2), e0192673. https://doi.org/10.1371/journal.pone.0192673

[2] Maroilley, T., Lemonnier, G., Lecardonnel, J., Esquerré, D., Ramayo-Caldas, Y., Mercat, M. J., Rogel-Gaillard, C., & Estellé, J. (2017). Deciphering the genetic regulation of peripheral blood transcriptome in pigs through expression genome-wide association study and allele-specific expression analysis. BMC genomics, 18(1), 967. https://doi.org/10.1186/s12864-017-4354-6

[3] Hui, L., Shuangshuang, G., Jianning, Y., & Zhendan, S. (2017). Systemic analysis of gene expression profiles in porcine granulosa cells during aging. Oncotarget, 8(57), 96588–96603. https://doi.org/10.18632/oncotarget.21731

[4] Velculescu, V. E., Zhang, L., Zhou, W., Vogelstein, J., Basrai, M. A., Bassett, D. E., Jr, Hieter, P., Vogelstein, B., & Kinzler, K. W. (1997). Characterization of the yeast transcriptome. Cell, 88(2), 243–251. https://doi.org/10.1016/s0092-8674(00)81845-0

[5] Skugor, A., Kjos, N. P., Sundaram, A., Mydland, L. T., Ånestad, R., Tauson, A. H., & Øverland, M. (2019). Effects of long-term feeding of rapeseed meal on skeletal muscle transcriptome, production efficiency and meat quality traits in Norwegian Landrace growing-finishing pigs. PloS one, 14(8), e0220441. https://doi.org/10.1371/journal.pone.0220441

[6] Guzek, D., Głąbska, D., Głąbski, K., & Wierzbicka, A. (2016). Influence of Duroc breed inclusion into Polish Landrace maternal line on pork meat quality traits. Anais da Academia Brasileira de Ciencias, 88(2), 1079–1088. https://doi.org/10.1590/0001-3765201620140679

[7] Alonso, V., Muela, E., Gutiérrez, B., Calanche, J. B., Roncalés, P., & Beltrán, J. A. (2015). The inclusion of Duroc breed in maternal line affects pork quality and fatty acid profile. Meat science, 107, 49–56. https://doi.org/10.1016/j.meatsci.2015.04.011

[8] Franco, D., Vazquez, J. A., & Lorenzo, J. M. (2014). Growth performance, carcass and meat quality of the Celta pig crossbred with Duroc and Landrance genotypes. Meat science, 96(1), 195–202. https://doi.org/10.1016/j.meatsci.2013.06.024

[9] Touma, S., & Oyadomari, M. (2020). Comparison of growth performances, carcass characteristics, and meat qualities of Okinawan indigenous Agu pigs and crossbred pigs sired by Agu or Duroc boar. Animal science journal = Nihon chikusan Gakkaiho, 91(1), e13362. https://doi.org/10.1111/asj.13362

[10] Li, X. J., Zhou, J., Liu, L. Q., Qian, K., & Wang, C. L. (2016). Identification of genes in longissimus dorsi muscle differentially expressed between Wannanhua and Yorkshire pigs using RNA-sequencing. Animal genetics, 47(3), 324–333. https://doi.org/10.1111/age.12421

[11] Xu, J., Wang, C., Jin, E., Gu, Y., Li, S., & Li, Q. (2018). Identification of differentially expressed genes in longissimus dorsi muscle between Wei and Yorkshire pigs using RNA sequencing. Genes & genomics, 40(4), 413–421. https://doi.org/10.1007/s13258-017-0643-3

[12] Wang, Z., Li, Q., Chamba, Y., Zhang, B., Shang, P., Zhang, H., & Wu, C. (2015). Identification of Genes Related to Growth and Lipid Deposition from Transcriptome Profiles of Pig Muscle Tissue. PloS one, 10(10), e0141138. https://doi.org/10.1371/journal.pone.0141138

[13] Hou, X., Yang, Y., Zhu, S., Hua, C., Zhou, R., Mu, Y., Tang, Z., & Li, K. (2016). Comparison of skeletal muscle miRNA and mRNA profiles among three pig breeds. Molecular genetics and genomics : MGG, 291(2), 559–573. https://doi.org/10.1007/s00438-015-1126-3

[14] Dos Santos, É. R., Bridi, A. M., da Silva, C. A., Alfieri, A. A., Fritzen, J., Terto, D. K., & Correia, E. R. (2021). Gender effects on pork quality and calpain-1 and calpastatin gene expression in male pig muscle. Meat science, 172, 108366. https://doi.org/10.1016/j.meatsci.2020.108366

[15] Braña, D. V., Rojo-Gómez, G. A., Ellis, M., & Cuaron, J. A. (2013). Effect of gender (gilt and surgically and immunocastrated male) and ractopamine hydrochloride supplementation on growth performance, carcass, and pork quality characteristics of finishing pigs under commercial conditions. Journal of animal science, 91(12), 5894–5904. https://doi.org/10.2527/jas.2013-6545

[16] Cameron, N. D., Warriss, P. D., Porter, S. J., & Enser, M. B. (1990). Comparison of Duroc and British landrace pigs for meat a and eating quality. Meat science, 27(3), 227–247. https://doi.org/10.1016/0309-1740(90)90053-9

[17] Liu, H., Xi, Y., Liu, G., Zhao, Y., Li, J., & Lei, M. (2018). Comparative transcriptomic analysis of skeletal muscle tissue during prenatal stages in Tongcheng and Yorkshire pig using RNA-seq. Functional & integrative genomics, 18(2), 195–209. https://doi.org/10.1007/s10142-017-0584-6

[18] Xu, J., Wang, C., Jin, E., Gu, Y., Li, S., & Li, Q. (2018). Identification of differentially expressed genes in longissimus dorsi muscle between Wei and Yorkshire pigs using RNA sequencing. Genes & genomics, 40(4), 413–421. https://doi.org/10.1007/s13258-017-0643-3

[19] Shen, J., Hao, Z., Wang, J., Hu, J., Liu, X., Li, S., Ke, N., Song, Y., Lu, Y., Hu, L., Qiao, L., Wu, X., & Luo, Y. (2021). Comparative Transcriptome Profile Analysis of Longissimus dorsi Muscle Tissues From Two Goat Breeds With Different Meat Production Performance Using RNA-Seq. Frontiers in genetics, 11, 619399. https://doi.org/10.3389/fgene.2020.619399

[20] Fraser, L., Brym, P., Pareek, C. S., Mogielnicka-Brzozowska, M., Paukszto, Ł., Jastrzebski, J. P., Wasilewska-Sakowska, K., Mańkowska, A., Sobiech, P., & Żukowski, K. (2020). Transcriptome analysis of boar spermatozoa with different freezability using RNA-Seq. Theriogenology, 142, 400–413. https://doi.org/10.1016/j.theriogenology.2019.11.001

[21] SSuomalainen, L., Dunkel, L., Ketola, I., Eriksson, M., Erkkilä, K., Oksjoki, R., Taari, K., Heikinheimo, M., & Pentikäinen, V. (2004). Activator protein-1 in human male germ cell apoptosis. Molecular human reproduction, 10(10), 743–753. https://doi.org/10.1093/molehr/gah094

[22] Keller, J., Ringseis, R., Priebe, S., Guthke, R., Kluge, H., & Eder, K. (2011). Dietary L-carnitine alters gene expression in skeletal muscle of piglets. Molecular nutrition & food research, 55(3), 419–429. https://doi.org/10.1002/mnfr.201000293

[23] Havis, E., & Duprez, D. (2020). EGR1 Transcription Factor is a Multifaceted Regulator of Matrix Production in Tendons and Other Connective Tissues. International journal of molecular sciences, 21(5), 1664. https://doi.org/10.3390/ijms21051664

[24] Cao, X. M., Guy, G. R., Sukhatme, V. P., & Tan, Y. H. (1992). Regulation of the Egr-1 gene by tumor necrosis factor and interferons in primary human fibroblasts. The Journal of biological chemistry, 267(2), 1345–1349.https://doi.org/10.1016/S0021-9258(18)48437-2

[25] Rockel, J. S., Bernier, S. M., & Leask, A. (2009). Egr-1 inhibits the expression of extracellular matrix genes in chondrocytes by TNFalpha-induced MEK/ERK signalling. Arthritis research & therapy, 11(1), R8. https://doi.org/10.1186/ar2595

[26] Aouadi, M., Laurent, K., Prot, M., Le Marchand-Brustel, Y., Binétruy, B., & Bost, F. (2006). Inhibition of p38MAPK increases adipogenesis from embryonic to adult stages. Diabetes, 55(2), 281–289. https://doi.org/10.2337/diabetes.55.02.06.db05-0963

[27] Zhang, D., Wu, W., Huang, X., Xu, K., Zheng, C., & Zhang, J. (2021). Comparative analysis of gene expression profiles in differentiated subcutaneous adipocytes between Jiaxing Black and Large White pigs. BMC genomics, 22(1), 61. https://doi.org/10.1186/s12864-020-07361-9

[28] Mohtar, O., Ozdemir, C., Roy, D., Shantaram, D., Emili, A., & Kandror, K. V. (2019). Egr1 mediates the effect of insulin on leptin transcription in adipocytes. The Journal of biological chemistry, 294(15), 5784–5789. https://doi.org/10.1074/jbc.AC119.007855

[29] Wang, Y., Ma, C., Sun, Y., Li, Y., Kang, L., & Jiang, Y. (2017). Dynamic transcriptome and DNA methylome analyses on longissimus dorsi to identify genes underlying intramuscular fat content in pigs. BMC genomics, 18(1), 780. https://doi.org/10.1186/s12864-017-4201-9

[30] Myers, M. J., Shepherd, D. L., Durr, A. J., Stanton, D. S., Mohamed, J. S., Hollander, J. M., & Alway, S. E. (2019). The role of SIRT1 in skeletal muscle function and repair of older mice. Journal of cachexia, sarcopenia and muscle, 10(4), 929–949. https://doi.org/10.1002/jcsm.12437

[31] Ding, R. B., Bao, J., & Deng, C. X. (2017). Emerging roles of SIRT1 in fatty liver diseases. International journal of biological sciences, 13(7), 852–867. https://doi.org/10.7150/ijbs.19370

[32] Azzu, V., & Brand, M. D. (2010). The on-off switches of the mitochondrial uncoupling proteins. Trends in biochemical sciences, 35(5), 298–307. https://doi.org/10.1016/j.tibs.2009.11.001

[33] Musa, C. V., Mancini, A., Alfieri, A., Labruna, G., Valerio, G., Franzese, A., Pasanisi, F., Licenziati, M. R., Sacchetti, L., & Buono, P. (2012). Four novel UCP3 gene variants associated with childhood obesity: effect on fatty acid oxidation and on prevention of triglyceride storage. International journal of obesity (2005), 36(2), 207–217. https://doi.org/10.1038/ijo.2011.81

[34] Huang, W., Zhang, X., Li, A., Xie, L., & Miao, X. (2017). Differential regulation of mRNAs and lncRNAs related to lipid metabolism in two pig breeds. Oncotarget, 8(50), 87539–87553. https://doi.org/10.18632/oncotarget.20978

